# Detection of plasmid contigs in draft genome assemblies using customized Kraken databases

**DOI:** 10.1101/2020.11.29.402966

**Authors:** Ryota Gomi, Kelly L. Wyres, Kathryn E. Holt

## Abstract

Plasmids play an important role in bacterial evolution and mediate horizontal transfer of genes including virulence and antimicrobial resistance genes. Although short-read sequencing technologies have enabled large-scale bacterial genomics, the resulting draft genome assemblies are often fragmented into hundreds of discrete contigs. Several tools and approaches have been developed to identify plasmid sequences in such assemblies, but require trade-off between sensitivity and specificity. Here we propose using the Kraken classifier, together with a custom Kraken database comprising known chromosomal and plasmid sequences of *Klebsiella pneumoniae* species complex (KpSC), to identify plasmid-derived contigs in draft assemblies. We assessed performance using Illumina-based draft genome assemblies for 82 KpSC isolates, for which complete genomes were available to supply ground truth. When benchmarked against five other classifiers (Centrifuge, RFPlasmid, mlplasmids, PlaScope, and Platon), Kraken showed balanced performance in terms of overall sensitivity and specificity (90.8% and 99.4%, respectively for contig count; 96.5% and >99.9%, respectively for cumulative contig length), and the highest accuracy (96.8% vs 91.8%-96.6% for contig count; 99.8% vs 99.0%-99.7% for cumulative contig length), and F1-score (94.5% vs 84.5%-94.1%, for contig count; 98.0% vs 88.9%-96.7% for cumulative contig length). Kraken also achieved consistent performance across our genome collection. Furthermore, we demonstrate that expanding the Kraken database with additional known chromosomal and plasmid sequences can further improve classification performance. Although we have focused here on the KpSC, this methodology could easily be applied to other species with a sufficient number of completed genomes.

**IMPACT STATEMENT:** The assembly of bacterial genomes using short-read data often results in hundreds of discrete contigs due to the presence of repeat sequences in those genomes. Separating plasmid contigs from chromosomal contigs in such assemblies is required, e.g., to assess the mobility of antimicrobial resistance genes. Although several tools have been developed for that purpose, they often suffer from low sensitivity or specificity. Here, we propose that the Kraken classifier coupled with a custom Kraken database comprising plasmid-free chromosomal sequences and complete plasmid sequences can be used for detection of plasmid contigs in draft genome assemblies. We showed that Kraken achieved balanced and higher performance compared with other methods (Centrifuge, RFPlasmid, mlplasmids, PlaScope, and Platon). We therefore consider that the Kraken classifier can be the best option for predicting the origin of contigs for species with a suitable number of completed chromosomal and plasmid sequences.

**DATA SUMMARY:** **Table S1**: Complete chromosomes used for creating the base Kraken database. Plasmid-free chromosomal sequences and complete plasmid sequences used for creating the base Kraken database are also available via Figshare at https://doi.org/10.6084/m9.figshare.13289564.

**Table S2**: Sequence data used for benchmarking. Draft assemblies of these 82 KpSC strains are available via Figshare at https://doi.org/10.6084/m9.figshare.13553432. The corresponding sequence read files and complete genomes were deposited in the NCBI SRA and GenBank under BioProjects PRJEB6891, PRJNA351909, PRJNA486877, and PRJNA646837 (individual BioSample IDs listed in Table S2).

Kraken output files are available via Figshare at https://doi.org/10.6084/m9.figshare.13553789.

## INTRODUCTION

Plasmids play an important role in horizontal genetic exchange and facilitate the movement and spread of virulence and antibiotic resistance genes.^1^ Accurate detection and analysis of these sequences is therefore a key component in understanding the epidemiology and evolution of bacterial pathogens, and can support genomic surveillance strategies. Previously, plasmids were extracted from bacterial cells, and subjected to targeted amplicon-based DNA sequencing for detailed characterization; however, these investigations were very labour-intensive, limiting the scale to which they could be applied. The advent of high-throughput sequencing has seen a transition from these labour-intensive approaches to massively-parallel sequencing of total genomic DNA, capturing both chromosome and plasmid sequences. The most widely used approach is Illumina sequencing, which provides highly accurate and economical short-read sequences. Long-read technologies such as those offered by Oxford Nanopore Technologies and Pacific Biosciences are increasingly used in bacterial genomics, although their use remains comparatively limited due to accuracy and/or cost constraints.

Due to the presence of repeat sequences, *de novo* assembly of bacterial genomes from short reads results in fragmented assemblies, generally comprising hundreds of contigs of unknown origins (chromosome or plasmid).^2^ Recently, several tools have been developed to identify plasmid sequences in such assemblies (or metagenomic sequence data), namely cBar,^3^ PlasFlow,^4^ Platon^5^, plasmidSPAdes,^6^ RFPlasmid,^7^ mlplasmids,^8^ and PlaScope.^9^ These tools can be divided into taxon-independent approaches (cBar, PlasFlow, Platon, plasmidSPAdes) and taxon-dependent approaches (mlplasmids, PlaScope) with the exception of RFPlasmid, which includes both taxon-dependent and taxon-independent models (**Table 1**). Taxon-dependent approaches have been reported to outperform taxon-independent approaches for their target taxon^7-9^ with the exception that Platon was reported by its developers to outperform PlaScope on 21 *Escherichia coli* genomes.^5^ Despite their advantages, each taxon-dependent tool has its own limitations. Currently, mlplasmids supports only three species (*Enterococcus faecium, E. coli, Klebsiella pneumoniae*), and it does not have functions that allow users to create new models for unsupported species (nor to train extended models using new data). RFPlasmid supports 17 taxa and also has a species-agnostic model for unsupported species, but the species-agnostic model was reported to show inferior performance compared with species specific models.^7^ PlaScope supports only two species (*E. coli* and *Klebsiella spp*.), and it does not accept assemblies generated using tools other than SPAdes,^10^ because it relies on parameters recorded in the headers of SPAdes-formatted fasta files to filter contigs. However, PlaScope is based on the Centrifuge classifier (a metagenomic classifier which classifies sequences based on exact matches against a database)^11^ and allows users to create their own Centrifuge databases for unsupported species, which makes the tool more flexible. Here, we propose that this concept can be improved by (i) using Kraken,^12^ an alternative metagenomic classifier based on the exact alignment of k-mers which also allows users to create their own databases and has been reported to provide higher precision than Centrifuge;^11^ (ii) refining the database by excluding chromosomal sequences containing integrated plasmid sequences.

**Table 1.** Characteristics of plasmid prediction tools.

| Tool | Classification features | Classification approach / method | Models / databases | Reference |
| --- | --- | --- | --- | --- |
| cBar | Pentamer frequencies | Sequential minimal optimization | Taxon-independent model | Zhou *et al.*<sup>3</sup> |
| PlasFlow | Kmer frequencies | Neural network | Taxon-independent model | Krawczyk *et al.*<sup>4</sup> |
| Platon | Chromosomal and plasmid marker protein sequences and other features (e.g. contig length) | Calculation of a mean replicon distribution score and higher-level contig characterizations | Taxon-independent database | Schwengers *et al.*<sup>5</sup> |
| plasmidSPAdes | Read coverage of contigs and structural features of the assembly graph | Assemble plasmids from whole genome sequencing data. | - | Antipov *et al.*<sup>6</sup> |
| RFPlasmid | Pentamer frequencies and chromosomal and plasmid marker genes | Random Forest | Taxon-specific models for 17 taxa and a taxon-independent model available | van der Graaf-van Bloois *et al.*<sup>7</sup> |
| mlplasmids | Pentamer frequencies | Support-vector machine | Three taxon-specific models available | Arredondo-Alonso *et al.*<sup>8</sup> |
| PlaScope | Exact matches to a database of known sequences | Centrifuge | Two taxon-specific databases available (novel databases can be created by the user) | Royer *et al.*<sup>9</sup> |

In this study, we first prepared a dataset comprising completed plasmids and chromosomes which are free from chromosomally integrated plasmids. We selected *Klebsiella pnemoniae* and closely related species (*K. quasipneumoniae, K. variicola*, and *K. quasivariicola*) within the *K. pneumoniae* species complex (KpSC) as study organisms because (i) *K. pneumoniae* has a higher plasmid burden than other Gram-negative opportunistic pathogens;^13^ (ii) these plasmids are of particular interest in genomic studies as they are frequently the location of functionally important genes involved in antimicrobial resistance and virulence;^14^ and (iii) the taxon-limited tools mlplasmids and PlaScope both support *K. pneumoniae*, making comparison with these high-performing alternatives straightforward. We used the same dataset to create custom databases for Kraken and Centrifuge; and benchmarked performance of these against RFPlasmid (using a taxon-dependent model), mlplasmids, PlaScope, and Platon. We chose RFPlasmid, mlplasmids, and PlaScope because they employ taxon-dependent models that have generally been shown to outperform other approaches, and Platon because it reportedly outperformed PlaScope.^5^

## METHODS

### Creation of Kraken and Centrifuge databases

An initial dataset was prepared by using the following component sequences: (i) complete chromosomes for *K. pneumoniae* (n=271), *K. quasipneumoniae* (n=13), *K. variicola* (n=14), and *K. quasivariicola* (n=1) downloaded from NCBI GenBank (**Table S1**); (ii) a publicly available dataset of complete Enterobacteriaceae plasmids (n=2097) reported previously.^15^ Plasmids can occasionally become integrated into chromosomes,^16^ or misassembled into chromosomal contigs. Inclusion of chromosomes with integrated plasmid sequences in databases could lead to erroneously classifying plasmid-derived contigs as chromosomal, hence chromosomal sequences containing integrated plasmid sequences were identified and removed by screening against known Enterobacteriaceae plasmid replicon markers using PlasmidFinder.^17^ The resulting dataset of plasmid-free chromosomal sequences and complete plasmid sequences was used to construct a Kraken database (hereafter referred as the ‘base’ database) using a taxonomic tree shown in **Figure S1** (also see **Supplementary Methods** for the commands used). A Centrifuge database was also created using the same dataset and the same taxonomic tree.

### Sequence data used for benchmarking

Complete genomes for clinical KpSC isolates (n=82) were used for benchmarking the performance of plasmid classifiers (**Table S2**). These isolates were sequenced via Illumina (short-read) and Oxford Nanopore Technologies (ONT, long-read) platforms as previously described.^16, 18-20^ To generate short-read only draft assemblies for testing as input to the classifiers, Illumina reads were first trimmed using Trim Galore (v0.5.0) (https://www.bioinformatics.babraham.ac.uk/projects/trim_galore/) and assembled using SPAdes (v3.13.1) with ╌careful and ╌only-assembler options.^10^ Contigs shorter than 1,000 bp or with a SPAdes contig coverage lower than 2x were discarded. In order to establish ‘ground truth’ classification of contigs with which to assess accuracy of classifier outputs, we completely resolved the genomes into circularized chromosome and plasmid sequences by assembling the ONT reads together with Illumina reads using Unicycler (v0.4.7).^21^ Genomes that did not completely resolve automatically using Unicycler were manually resolved as described previously.^19^

Contigs in each draft (short-read only) assembly were labeled as plasmid-derived or chromosome-derived by mapping to the corresponding complete genome for the same isolate, using minimap2 (v2.14) with
 −cx asm5 option.^22^ Contigs that either (i) mapped to both chromosomes and plasmids, (ii) failed to map at all or (iii) yielded no significant alignment (i.e., no alignment block with length ≥50% of the contig), were discarded (median two contigs and 3,371 bp per genome; see **Table S2**).

Chromosomal multi-locus sequence types (STs) were determined using Kleborate (v0.4.0-beta) (https://github.com/katholt/Kleborate) to assess the diversity of the genomes used for benchmarking.

### Performance benchmarks

Contigs from short-read only draft assemblies were classified as plasmid or chromosomal using Kraken, Centrifuge, RFPlasmid, mlplasmids, PlaScope, and Platon (see **Supplementary Method**s for the commands used). Kraken (v1.1.1) was run with default settings using the base database created above. Contigs assigned taxonomy ID 0 (not in the database) or 1 (root) were considered unclassified (**Figure S1**). Centrifuge (v1.0.4) was run using the database created above (centrifuge −f − p 10 ╌reorder −x database −U sequence.fasta −k 1 ╌report-file sequence_summary.txt −S sequence_output.txt). Contigs assigned taxonomy ID 0 or 1 were considered unclassified. RFPlasmid (v0.0.9) was run using the Enterobacteriaceae model. mlplasmids (v1.0.0) was run using the following settings: full_output = TRUE, species = “Klebsiella pneumoniae". PlaScope (v1.3.1) was run with mode 2 using a *Klebsiella* database provided by the developers. Platon (v1.4.0) was run with default settings.

To calculate performance statistics, the category plasmid was considered as the positive-class, and the category chromosome was considered as the negative-class. Moreover, Kraken, Centrifuge, and PlaScope may assign contigs as unclassified for ambiguous results. Prediction for each contig was compared with the ground truth determined by minimap2 mapping to completely resolved hybrid assemblies as described above, and results assigned to categories: true positive (TP) (predicted as plasmid-derived and mapped to true plasmids), true negative (TN) (predicted as chromosome-derived/unclassified and mapped to true chromosome), false positive (FP) (predicted as plasmid-derived but mapped to true chromosome), or false negative (FN) (predicted as chromosome-derived/unclassified but mapped to true plasmid). Sensitivity [TP/(TP+FN)], specificity [TN/(TN+FP)], precision [TP/(TP+FP)], negative predictive value [TN/(TN+FN)], accuracy [(TP+TN)/(TP+TN+FP+FN)], and F1-score [2×sensitivity×precision/(sensitivity+precision)] were calculated. These metrics were calculated in terms of raw contig counts (contig-wise) as well as cumulative contig length (nucleotide-wise) to account for genome-specific differences in contig length distributions.

### Assessing applicability of Kraken for classification of contigs with antimicrobial resistance genes

Antimicrobial resistance genes were detected in each draft assembly comprising the benchmarking dataset using Abricate (v1.0.1, https://github.com/tseemann/abricate). Abricate was run using the Resfinder database^23^ with the following thresholds: a minimum DNA identity of 95% and a minimum DNA coverage of 80%. Kraken predictions and assigned categories were retrieved for contigs in which antimicrobial resistance genes were detected.

### Assessing improvements in performance achieved by expanding the Kraken database

In order to assess the performance improvement achieved by expanding the base Kraken database we used a 3-fold cross-validation approach whereby the 82 benchmarking genomes were randomly divided into three groups: A (n=27), B (n=27), and C (n=28). Completed chromosomes and plasmids from groups A and B were added to the base database set to create an expanded base+A+B Kraken database with which to assess the classification of draft contigs from group C. This procedure was repeated using draft genomes from group A as a test set against an expanded base+B+C database, and draft genomes from group B as a test set against an expanded base+A+C database. Sensitivity, specificity, precision, negative predictive value, accuracy, and F1-score were calculated for each repetition.

## RESULTS AND DISCUSSION

### Benchmarking dataset characteristics

The benchmarking genome set was highly diverse, comprising 43 STs of *K. pneumoniae* (n=69), 1 ST of *K. quasipneumoniae* (n=1), and 8 STs of *K. variicola* (n=12). The draft assemblies of these genomes had a median of 63 contigs and a median N50 of 214,404 bp after discarding contigs shorter than 1,000 bp or with a SPAdes contig coverage lower than 2x. Hybrid assemblies revealed that these strains carried 0 to 9 plasmids (median 3, interquartile range 5-2; see **Table S2**). A total of 301 completed plasmids were assembled, and these sequences ranged from 1,240 bp to 430,829 bp (median 53,776 bp, also see **Figure S2** for the length distribution of the completed plasmids).

### Kraken-based contig classification outperforms existing approaches

Kraken, Centrifuge, RFPlasmid, mlplasmids, PlaScope, and Platon were run on the 82 KpSC genomes and aggregated metrics were calculated for benchmarking (**Table 2**). Performance metrics were also calculated for each individual genome (**Figure S3**) and the distributions are shown in **Figure 1**. All the tools performed well in terms of specificity, negative predictive values, and accuracy when contig length was considered as a unit (overall values >99.7%, >98.9% and >98.9%, respectively). However, metrics calculated for contig counts tended to be lower for all six tools, which indicates misclassification mainly occurred for short contigs for all the tools (**Figure S4**). Overall, Kraken showed the most balanced performance in terms of sensitivity (90.8% and 96.5% for aggregated contig counts and cumulative contig lengths, respectively; **Table 2**) and specificity (99.4% and >99.9% for aggregated contig counts and cumulative contig lengths, respectively; **Table 2**), and achieved the highest Youden Index (0.902 vs 0.736–0.897 for aggregated contig count; 0.965 vs 0.802–0.957 for cumulative contig length).^24^ Although mlplasmids performed better in terms of sensitivity and negative predictive values calculated for contig counts, this seems to be achieved at the cost of increased number of false positives (**Table S3**, median two contigs and 3,617 bp per genome). Similarly, Platon achieved the highest specificity (>99.9% for contig length and 99.4% for contig count) but showed the lowest sensitivity (80.2% for contig length and 74.2% for contig count). Finally, Kraken also achieved the highest accuracy (96.8% vs 91.8%-96.6% for aggregated contig count; 99.8% vs 99.0%-99.7% for cumulative contig length), and F1-score (94.5% vs 84.5%-94.1%, for aggregated contig count; 98.0% vs 88.9%-96.7% for cumulative contig length).

**Table 2.** Performance metrics for Kraken, Centrifuge, RFPlasmid, mlplasmids, PlaScope, and Platon on contigs from 82 KpSC genomes.

|  | Kraken | Centrifuge | RFPlasmid | mlplasmids | PlaScope | Platon |
| --- | --- | --- | --- | --- | --- | --- |
| **Sensitivity (true positive rate, %)** |  |  |  |  |  |  |
| Contig length | **96.507** | 94.565 | 94.198 | 95.987 | 87.199 | 80.215 |
| Contig count | 90.796 | 90.485 | 82.898 | **95.833** | 81.157 | 74.192 |
| **Specificity (true negative rate, %)** |  |  |  |  |  |  |
| Contig length | 99.973 | 99.941 | 99.859 | 99.727 | 99.958 | **99.984** |
| Contig count | 99.378 | 99.189 | 98.458 | 91.804 | 99.243 | **99.405** |
| **Precision (positive predictive value, %)** |  |  |  |  |  |  |
| Contig length | 99.472 | 98.851 | 97.289 | 94.965 | 99.120 | **99.638** |
| Contig count | **98.449** | 97.980 | 95.899 | 83.568 | 97.899 | 98.189 |
| **Negative predictive value (%)** |  |  |  |  |  |  |
| Contig length | **99.813** | 99.709 | 99.689 | 99.784 | 99.317 | 98.948 |
| Contig count | 96.128 | 95.995 | 92.976 | **98.064** | 92.372 | 89.853 |
| **Accuracy (%)** |  |  |  |  |  |  |
| Contig length | **99.796** | 99.667 | 99.570 | 99.536 | 99.308 | 98.977 |
| Contig count | **96.777** | 96.550 | 93.742 | 93.025 | 93.761 | 91.762 |
| **F1-score (%)** |  |  |  |  |  |  |
| Contig length | **97.967** | 96.660 | 95.719 | 95.474 | 92.778 | 88.878 |
| Contig count | **94.468** | 94.083 | 88.926 | 89.282 | 88.745 | 84.520 |

**Figure 1.**
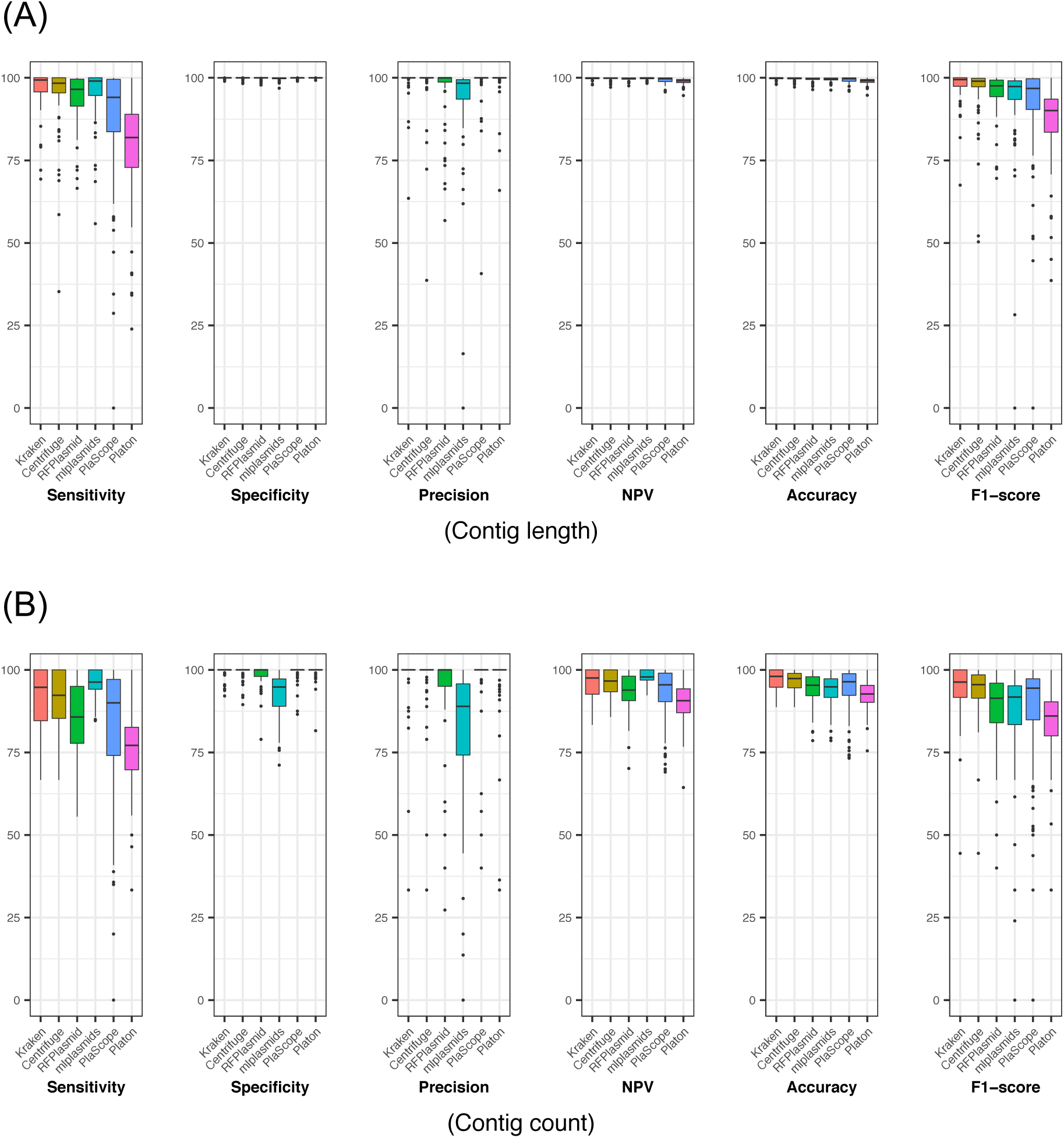
Performance metrics for Kraken, Centrifuge, RFPlasmid, mlplasmids, PlaScope, and Platon for each genome in terms of contig length (A) and contig count (B). NPV, negative predictive value.

We also note that there was variation in performance by genome, particularly for sensitivity, precision, and F1-score (**Figure 1, Figure S3, Table S4**). Both Kraken and Centrifuge achieved consistent performance, especially for metrics calculated for contig length (e.g., the interquartile ranges of F1-scores were 2.60% and 2.53% for Kraken and Centrifuge, respectively while the interquartile ranges of F1-scores for the other tools ranged from 5.03% to 9.99%, also see **Table S4**). Notably, the median values for Kraken were equal to or higher than those for Centrifuge for all metrics (**Table S4**), although the differences in the distributions were not considered statistically significant (**Table S5**). In contrast, the majority of comparisons of distributions between Kraken and the other approaches (n=38/48 and in 32/38 cases the median was higher for Kraken) were considered statistically significant (**Table S5**), commensurate with the general trend of broader distributions and/or lower values achieved for these approaches (**Figure 1**).

Notably, in our analysis sensitivity was higher for Centrifuge alone than PlaScope, which also uses the Centrifuge classifier internally. Chromosomes with integrated plasmid sequences were filtered out in our Centrifuge database but not in the Centrifuge database used in PlaScope, which seems to be the main reason for this difference: Inclusion of chromosomes with integrated plasmid sequences in our Centrifuge database reduced the number of true positives by 199 (from 1,455 to 1,256) and decreased sensitivity (**Table S6**), supporting this hypothesis. Minimap2 mapped 194 of those 199 contigs to chromosomally integrated plasmids (alignment block length ≥ 50% of the contig length).

Since mlplasmids was mainly designed for *K. pneumoniae*, we assessed the performance of mlplasmids excluding non-*K. pneumoniae* genomes (**Table S7**). As expected, performance of mlplasmids was slightly better for *K. pneumoniae* in most metrics (mean 0.261% increase for contig length and 0.966% increase for contig count), though precision remained relatively low (95.8 % for contig length and 86.1% for contig count). We also assessed the performance of mlplasmids by changing the probability threshold. By default, mlplasmids assigns each contig as plasmid-derived or chromosome-derived with a highest posterior probability. We changed this threshold (0.5) to 0.7, which is the value used for predicting the location of antibiotic-resistance genes in the original paper,^8^ and assessed the performance of mlplasmids by using the 82 KpSC genomes. **Table S8** shows that filtering out contigs based on a minimum posterior probability can lead to the increase of specificity (0.155% increase for contig length and 6.52% increase for contig count) and precision (2.65% increase for contig length and 11.8% increase for contig count), although sensitivity decreases (5.75% decrease for contig length and 17.0% decrease for contig count).

### Performance of Kraken for classification of contigs with antimicrobial resistance genes

A total of 641 resistance genes were detected in the 82 draft genome assemblies, but 11 of them were located on contigs that either (i) mapped to both chromosomes and plasmids, (ii) failed to map at all or (iii) yielded no significant alignment, and thus were not considered in the following analysis (see **Methods**). Kraken correctly predicted the locations of 568 genes (308 chromosome-derived and 260 plasmid-derived), while misclassified only 10 genes (seven chromosomal genes predicted as plasmid-derived and three plasmid genes predicted as chromosome-derived). The remaining 52 genes were on 38 contigs that could not be classified by Kraken (median 2,710 bp, range 1,227 bp to 6,981 bp). Kraken assigned taxonomy ID of 1 (root) to all 38 contigs, indicating those unclassified resistance genes are on mobile genetic elements that can be located both on chromosomes and plasmids. BLASTN search against Genbank database revealed all 38 contigs have hits with 100% query coverage and 100% identity for one or more chromosomes and one or more plasmid sequences.

### Impact of expanding the Kraken database

We assessed the performance improvement achieved by expanding the base Kraken database, which resulted in a marginal improvement to all the metrics except for sensitivity calculated for contig counts (0.002%-1.04% increase for contig length and 0.008%-0.534% increase for contig counts, **Table S9**) (p = 0.06 for McNemar’s test for differences in specificity)^25^. Compared with the classification results using the base database, 87 out of 5305 contigs (1.6%) were classified into different categories when the expanded databases were used (**Figure 2, Table S10**). Among them, 30 contigs were reclassified into the correct classes after expanding the database. Moreover, marked decrease was observed in the number of misclassified contigs, i.e. false positives+false negatives (classified), when the expanded databases were used (from 40 contigs to 8 contigs). On the other hand, the number of unclassified contigs increased by 39 after expanding the database due to the increase of contigs with taxonomy ID 1 (root). This implies using a larger number of genomes for creating a Kraken database can increase the number of mapped *k*-mers in the Kraken analysis, which can also increase the number of *k*-mers mapped both to chromosomes and plasmids and thus can lead to the increase of unclassified cases. Importantly, antimicrobial resistance genes were present in 22 contigs that were correctly classified as plasmid-derived using the base database but were assigned taxonomy ID 1 using the expanded databases, resulting in the decrease of the sensitivity for detection of resistance genes. All but one of those 22 contigs had hits with query coverage of 100% and percent identity of 100% against both chromosomal and plasmid sequences in Genbank database, again highlighting the difficulties of predicting the origins of contigs on mobile genetic elements.

**Figure 2.**
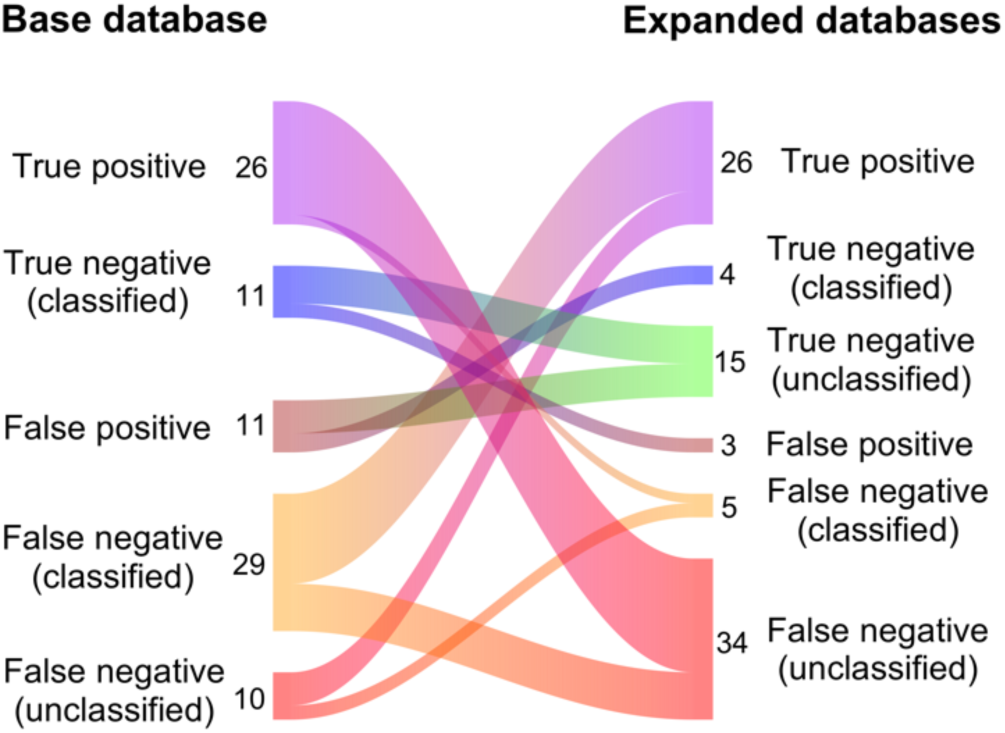
Classification results for 87 contigs that were classified into different categories after expanding the Kraken database. Contig number for each category is shown next to the diagram. Note that contigs classified into the same category are not shown in this figure (n = 5218 contigs). The Sankey diagram was created using the riverplot R package (v0.6).

## Conclusions

Here, we showed that the Kraken sequence classifier applied with a custom curated database can be used for detecting plasmid contigs in draft genome assemblies with high sensitivity (96.5% for contig length and 90.8% for contig count) and specificity (>99.9% for contig length and 99.4% for contig count). Notably, this approach was among the most consistent and balanced performers in comparison to tools designed specifically for this task (RFPlasmid, mlplasmids, PlaScope, and Platon), and in comparison to Centrifuge sequence classifier applied with the same database. This approach can be readily implemented for draft genome assemblies created with any assembly software because it does not require a specific fasta header format for input files (unlike PlaScope, which requires the SPAdes fasta header format for determination of contig depth). Moreover, performance can be further improved by expanding the Kraken database-a process that can be easily achieved through updating and rebuilding the database, although we note that this may also lead to the increase of unclassified contigs, particularly for those representing mobile genetic elements that can insert within both plasmids and chromosomes. In contrast, it is more difficult for users to modify the performance of other tools such as mlplasmids that require not only updating a database but re-training the classifier.

Although we tested the Kraken classifier only on KpSC, this methodology can be applied to other species for which a suitable number of completed chromosomal and plasmid sequences are available to build a database, and/or their close relatives. However, it may not be useful for species that have rarely been sequenced, though this is also the case for other taxon-dependent classifiers requiring pre-built databases or models. If an appropriate database were available, it is also possible that Kraken could be used for taxon-independent analyses and/or classification of plasmids from metagenomic assemblies, although further studies are needed to explore its accuracy for these use-cases. Nonetheless, for well-described species Kraken can be the best option to predict the origin of contigs from draft genome assemblies, to determine the location of antimicrobial resistance and virulence genes, and ultimately inform genomic surveillance studies.

### Funding information

This work was supported by the John Mung Program from Kyoto University, Japan (R.G.), JSPS KAKENHI (grant JP19K20461 to R.G.), the National Health and Medical Research Council of Australia (Investigator Grant APP1176192 to K.L.W), the Viertel Charitable Foundation of Australia (Senior Medical Research Fellowship to K.E.H.) and the Bill and Melinda Gates Foundation of Seattle (grant OPP1175797 to K.E.H.).

## Author contributions

Conceptualization, R.G., K.W., and K.E.H.; Methodology, R.G., K.W., and K.E.H.; Validation, R.G.; Writing – Original Draft Preparation, R.G.; Writing – Review and Editing, R.G., K.W., and K.E.H.; Supervision, K.E.H.

## Supplementary Figures

**Figure S1.**
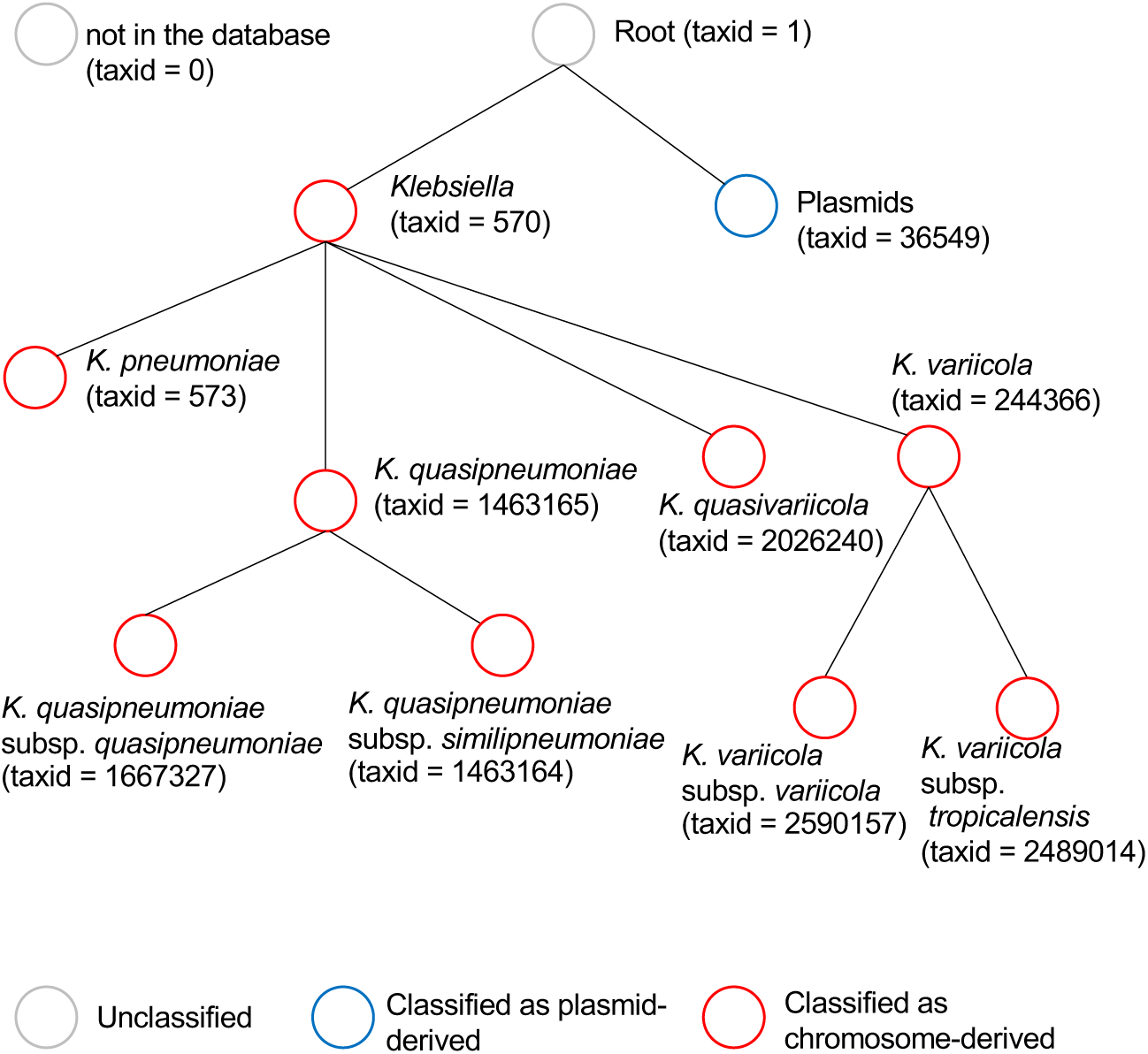
The taxonomic tree used in the Kraken analysis. This is a taxonomy tree from the NCBI taxonomy downloaded by “kraken-build ╌download-taxonomy ╌db database”. Although there are other taxa in the tree, they are not shown here because no chromosomal sequences other than *Klebsiella* spp. were included in the database. One of the taxonomic IDs in the tree was assigned to each contig. Contigs with a taxonomic ID 0 or 1 were labeled as unclassified, those with an ID 36549 were classified as plasmid-derived, and those with the remaining IDs were classified as chromosome-derived. Note that the taxon *Klebsiella variicola* subsp. *tropicalensis* has been renamed to *K. variicola* subsp. *tropica*, but has not yet been updated in NCBI (https://www.sciencedirect.com/science/article/pii/S0923250819300956).

**Figure S2.**
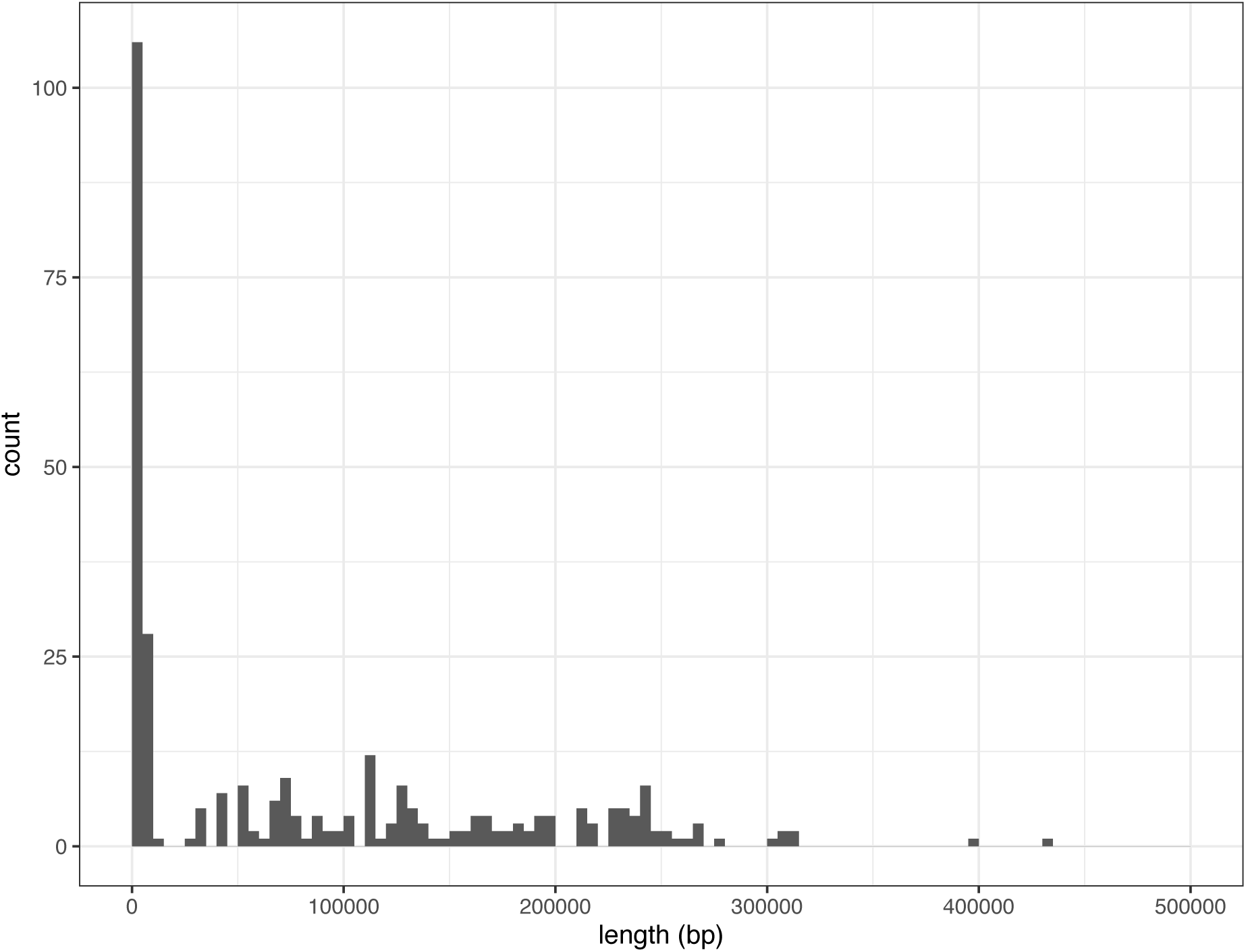
Length distribution of the completed plasmids (n=301).

**Figure S3.**
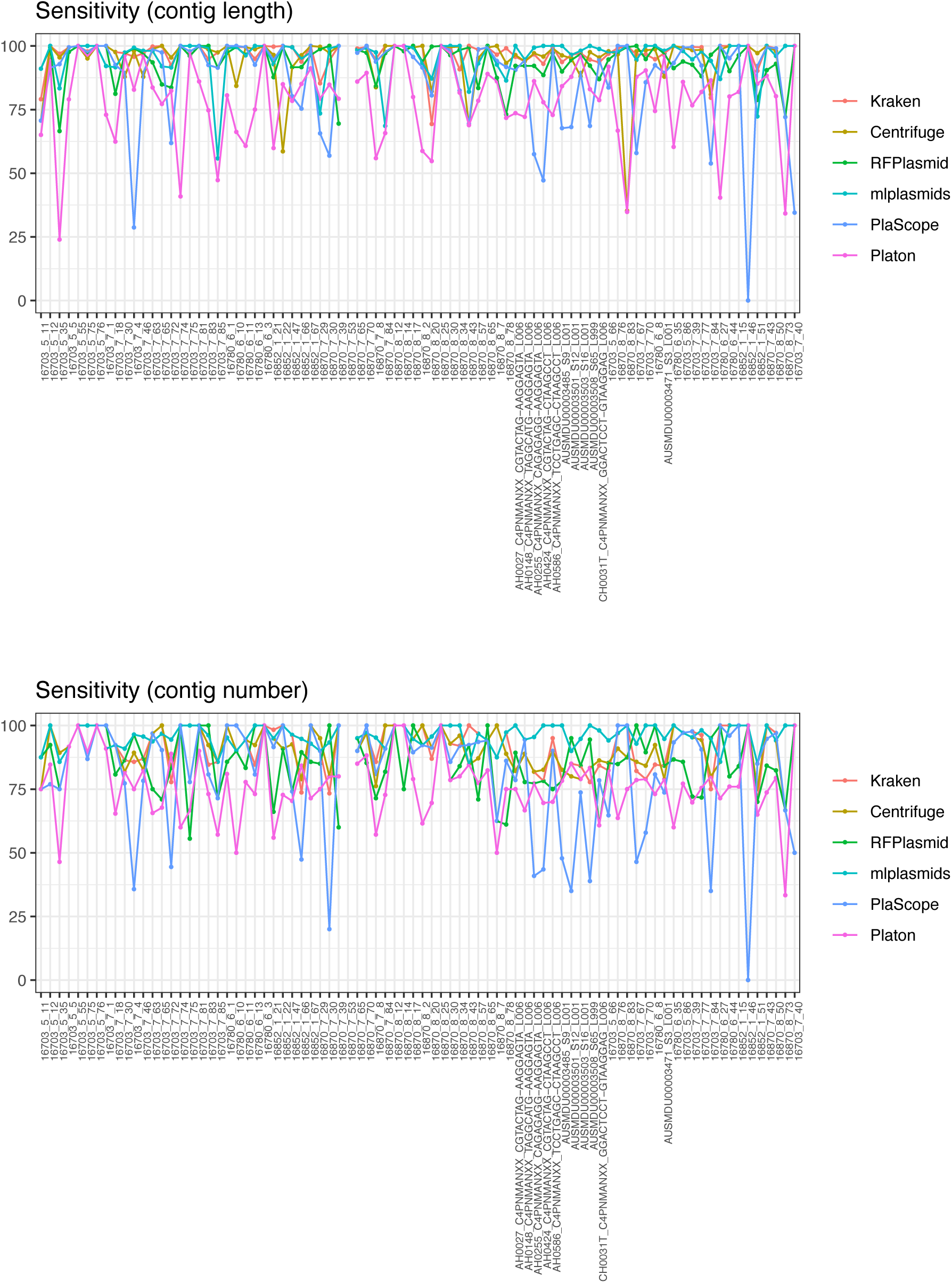
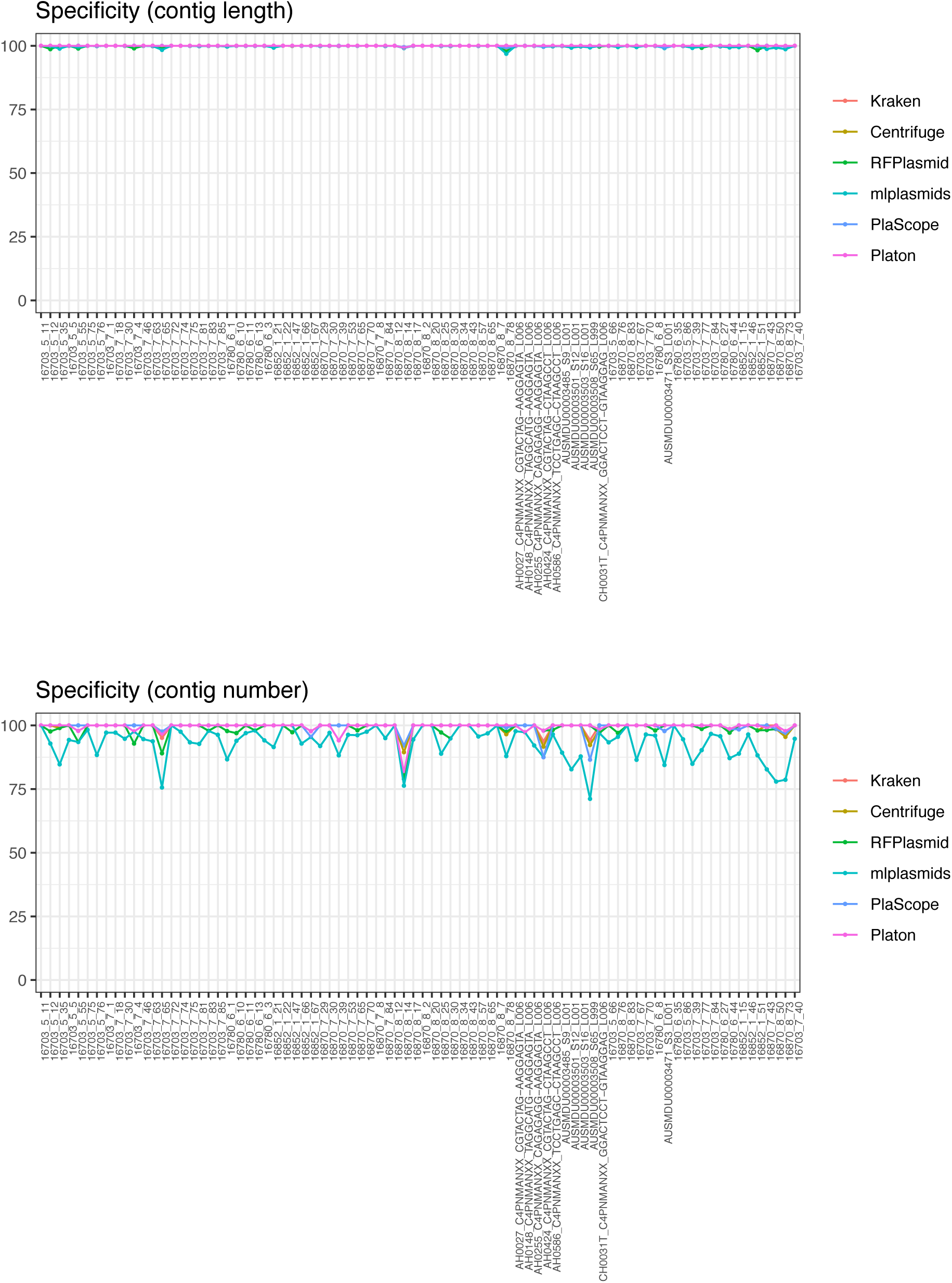
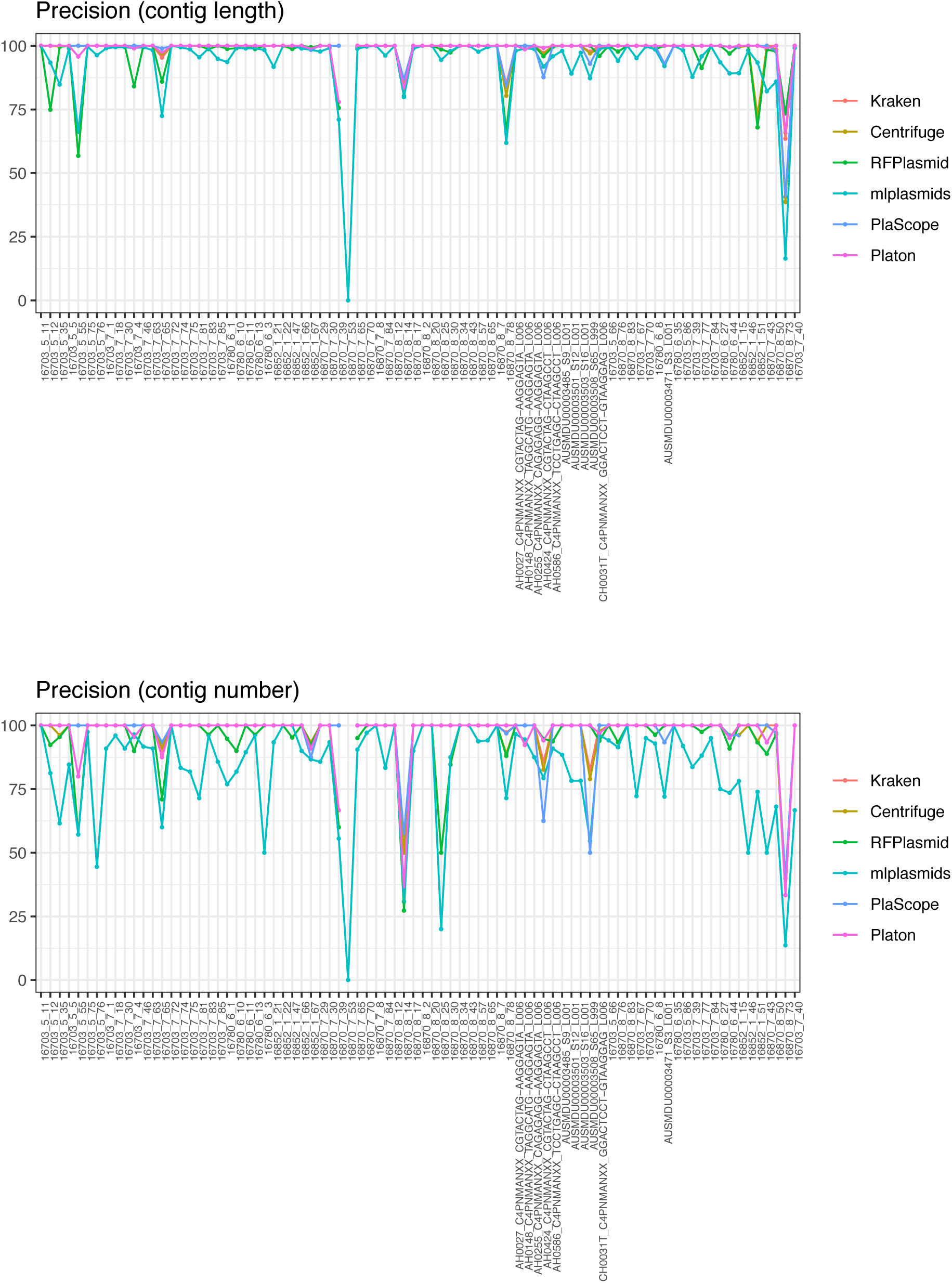
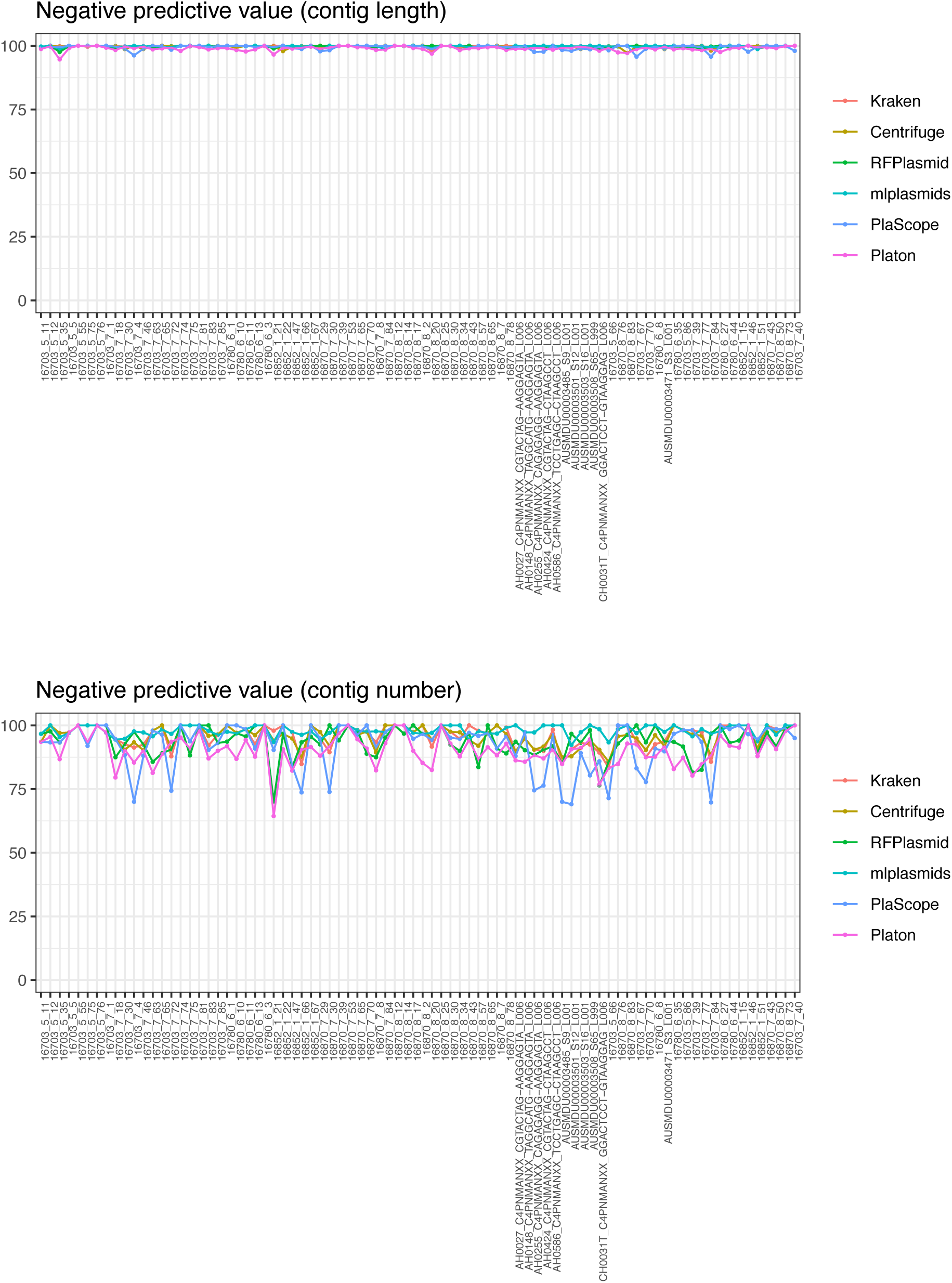
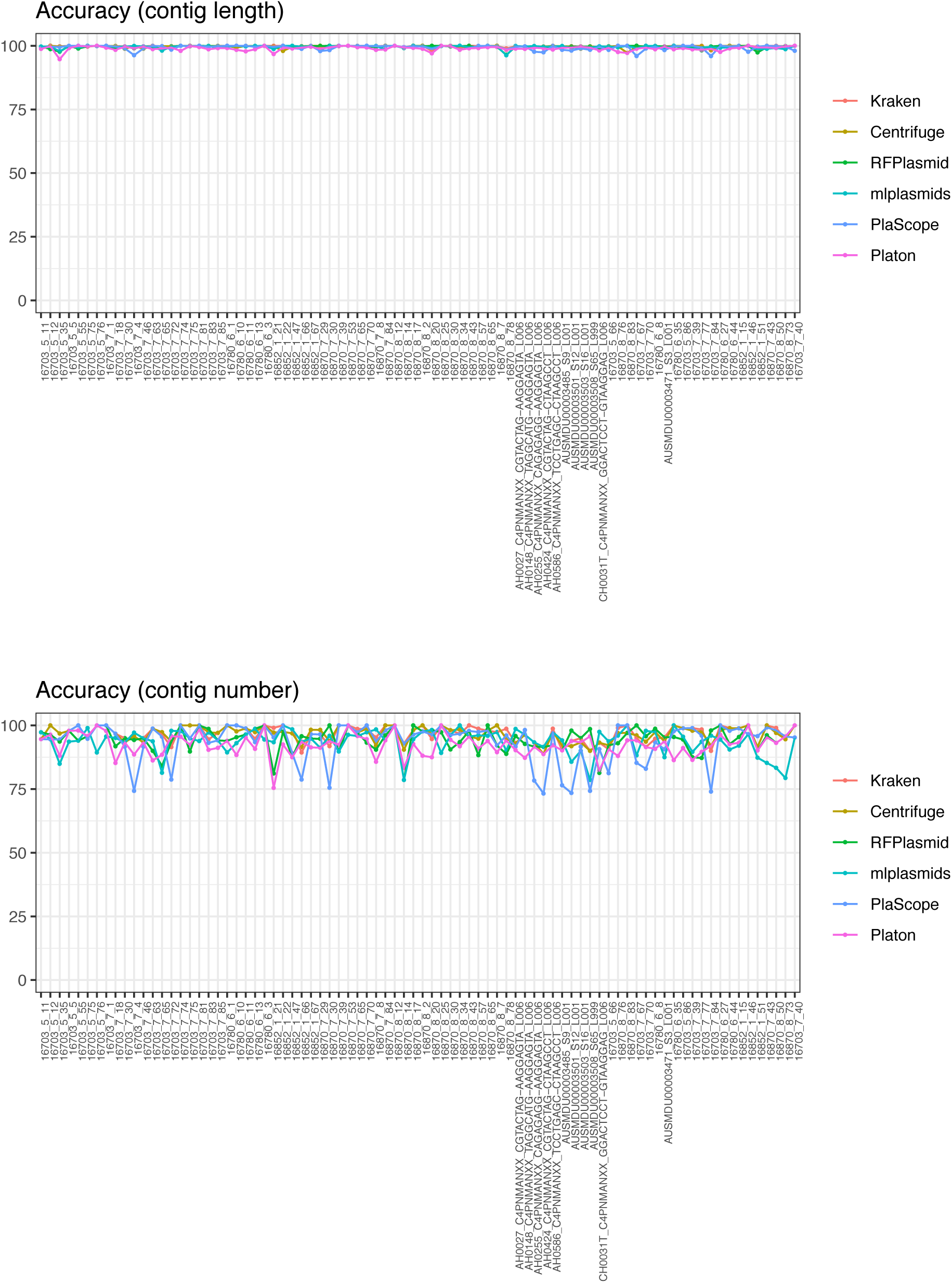
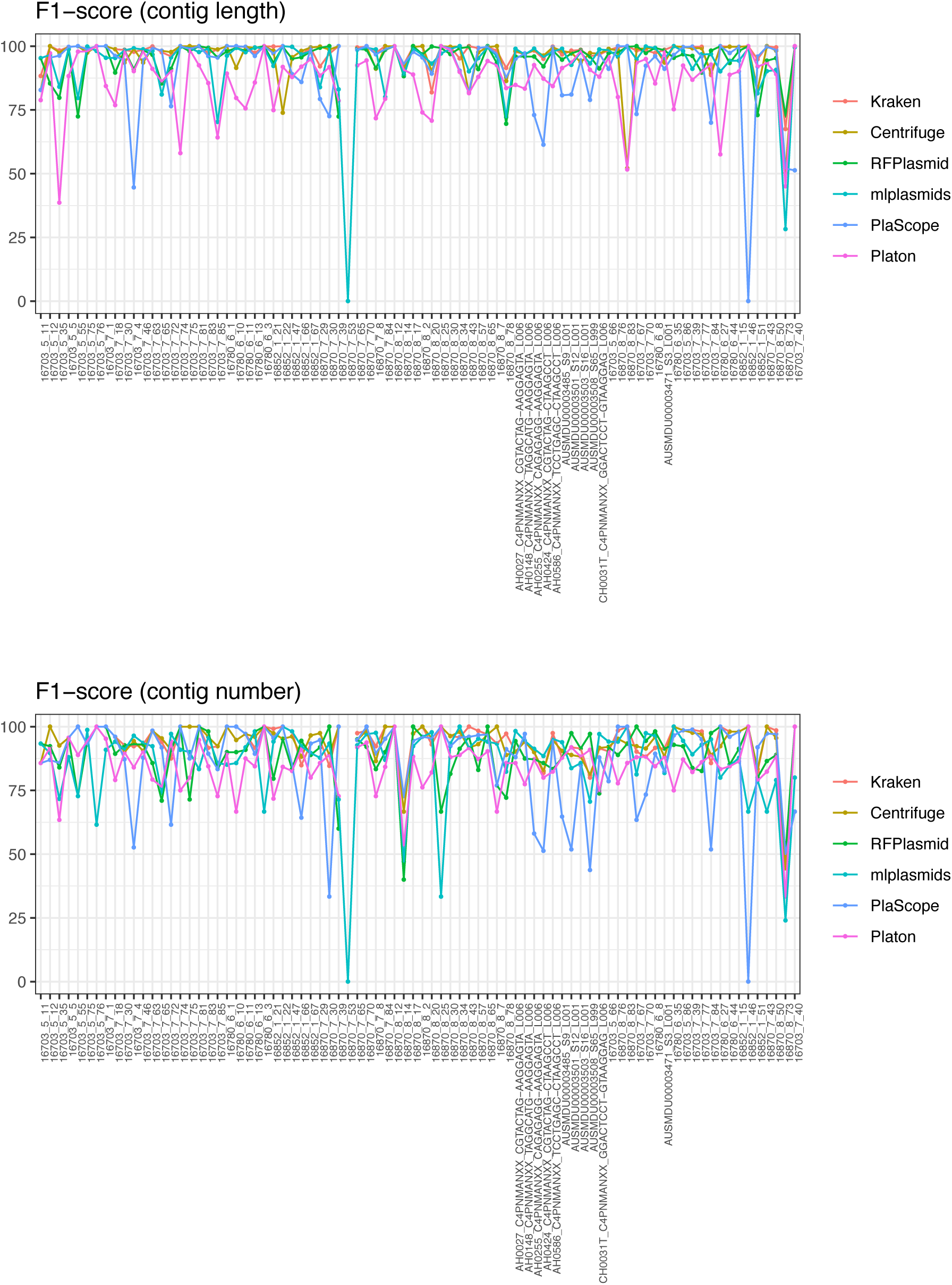
Performance metrics for Kraken, Centrifuge, RFPlasmid, mlplasmids, PlaScope, and Platon for each genome. Sensitivity, precision, and F1-score are not shown for 16870_7_53 except mlplasmids because this genome does not include plasmids and thus the true positive category is not applicable. (mlplasmids predicted one contig as plasmid-derived for this genome.)

**Figure S4.**
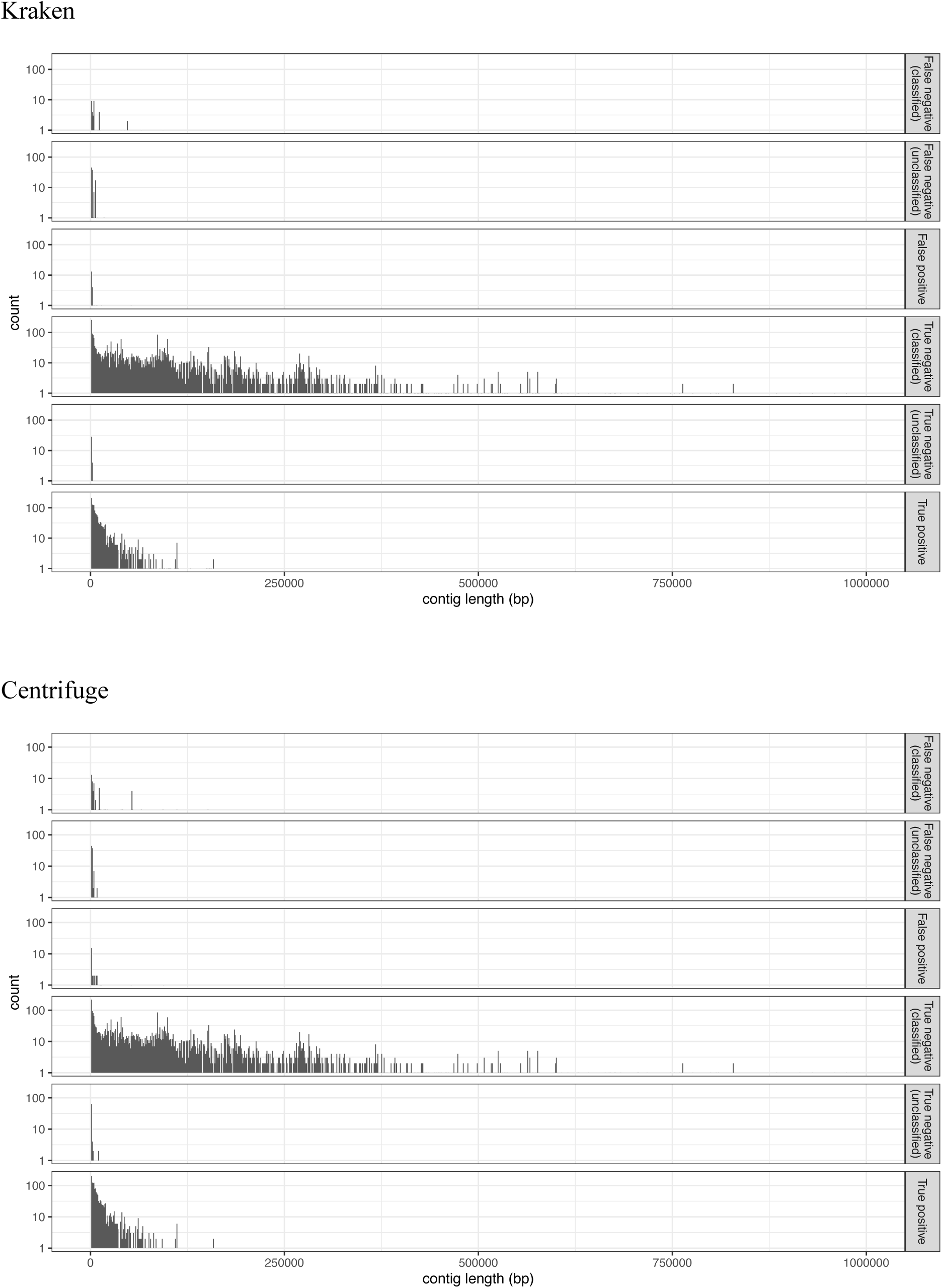
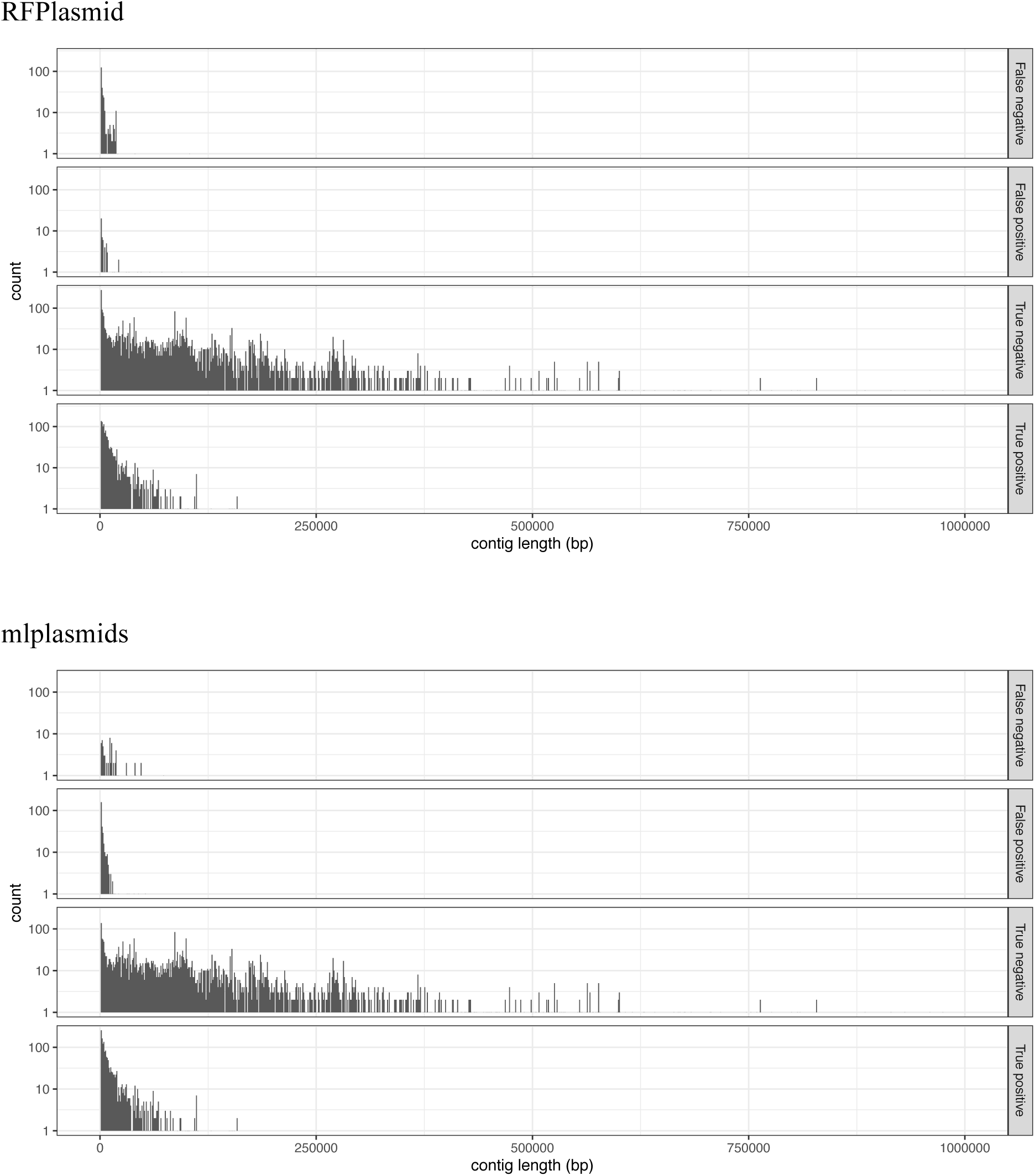
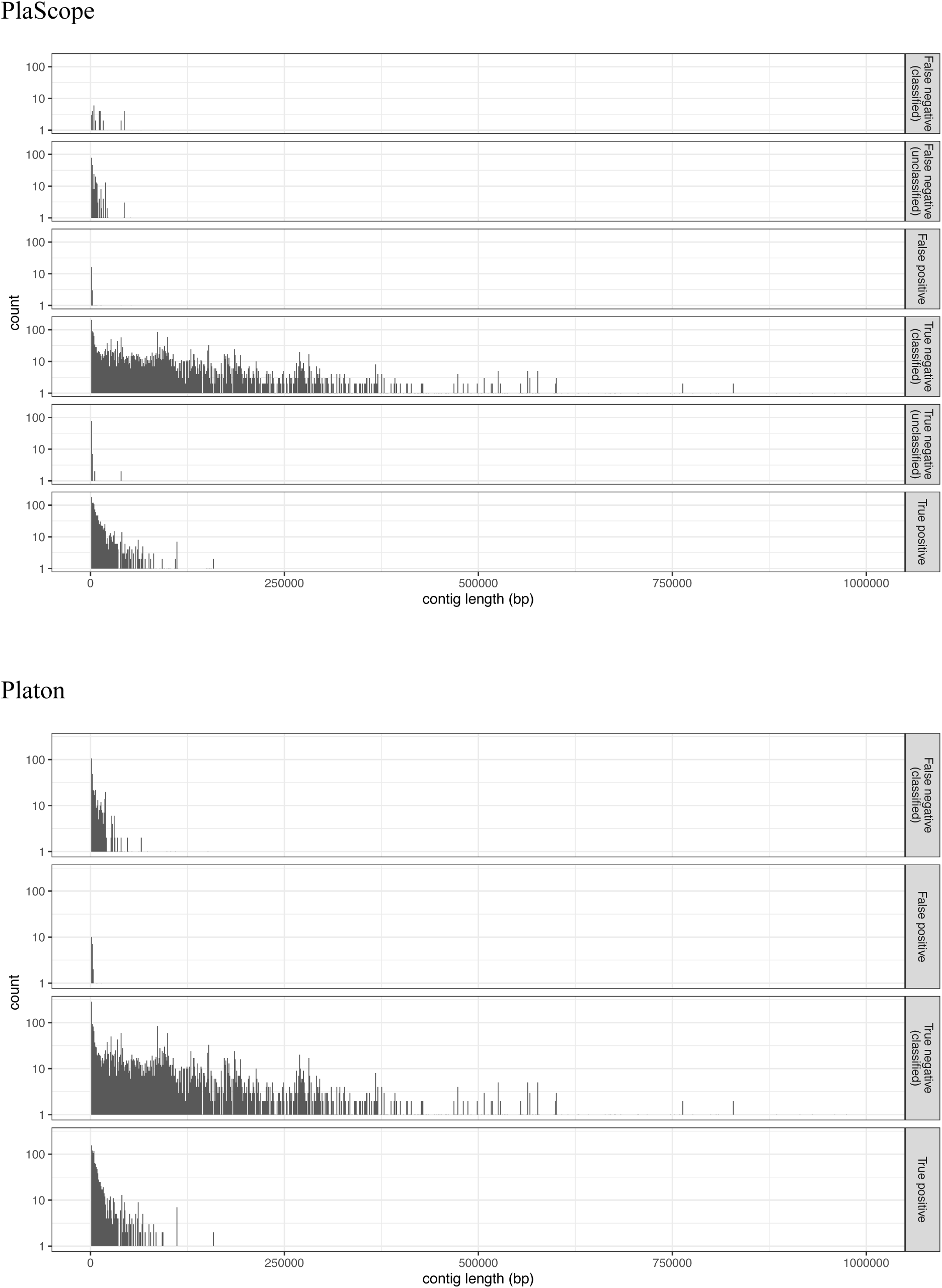
Relationships between contig length and classification categories. The y axes are on a log10 scale.

## Supplementary Tables

**Table S1.** See the excel file.

**Table S2.** See the excel file.

**Table S3.**
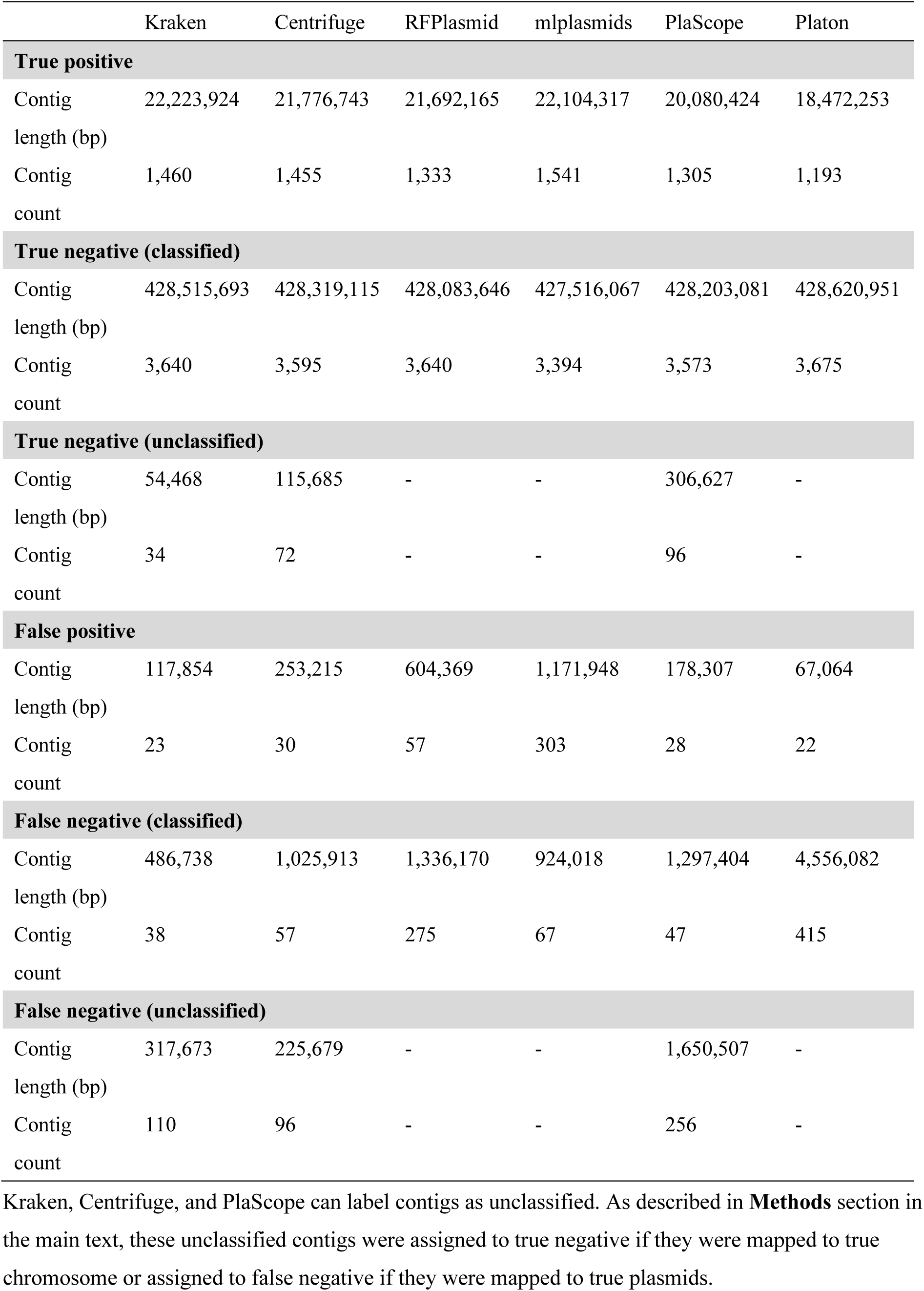
Total contig sizes and contig counts assigned to each category.

**Table S4.** Consistency of performance between genomes in terms of interquartile range (IQR). Median values are shown in parentheses. Bold font indicates the lowest IQR and highest median value(s) in each category.

|  | Kraken | Centrifuge | RFPlasmid | mlplasmids | PlaScope | Platon |
| --- | --- | --- | --- | --- | --- | --- |
| **IQR of sensitivity (%)** |  |  |  |  |  |  |
| Contig | **4.27** | 4.55 | 8.15 | 5.36 | 15.88 | 16.10 |
| length | **(99.32)** | (98.38) | (96.51) | (99.05) | (94.08) | (81.91) |
| Contig | 15.38 | 14.71 | 17.22 | **5.88** | 23.07 | 12.84 |
| count | (94.74) | (92.31) | (85.71) | **(96.30)** | (90.00) | (77.14) |
| **IQR of specificity (%)** |  |  |  |  |  |  |
| Contig | **0** | **0** | 0.06 | 0.27 | **0** | **0** |
| length | **(100)** | **(100)** | **(100)** | (99.93) | **(100)** | **(100)** |
| Contig | **0** | **0** | 1.99 | 8.27 | **0** | **0** |
| count | **(100)** | **(100)** | **(100)** | (94.80) | **(100)** | **(100)** |
| **IQR of precision (%)** |  |  |  |  |  |  |
| Contig | **0** | **0** | 1.32 | 5.91 | **0** | **0** |
| length | **(100)** | **(100)** | **(100)** | (98.39) | **(100)** | **(100)** |
| Contig | **0** | **0** | 5.00 | 21.57 | **0** | **0** |
| count | **(100)** | **(100)** | **(100)** | (88.97) | **(100)** | **(100)** |
| **IQR of negative predictive value (%)** |  |  |  |  |  |  |
| Contig | 0.27 | **0.26** | 0.42 | 0.27 | 1.10 | 0.93 |
| length | **(99.96)** | (99.91) | (99.82) | (99.95) | (99.76) | (99.07) |
| Contig | 7.38 | 6.61 | 7.41 | **3.10** | 8.65 | 7.25 |
| count | (97.53) | (96.64) | (93.89) | **(97.85)** | (95.48) | (90.65) |
| **IQR of accuracy (%)** |  |  |  |  |  |  |
| Contig | **0.28** | 0.29 | 0.42 | 0.55 | 1.01 | 0.87 |
| length | **(99.95)** | (99.90) | (99.70) | (99.75) | (99.76) | (99.10) |
| Contig | 5.26 | **4.36** | 5.74 | 5.61 | 6.56 | 5.17 |
| count | **(98.04)** | (97.36) | (95.33) | (94.85) | (96.40) | (92.68) |
| **IQR of F1-score (%)** |  |  |  |  |  |  |
| Contig | 2.60 | **2.53** | 5.03 | 5.63 | 9.33 | 9.99 |
| length | **(99.48)** | (99.01) | (97.60) | (97.37) | (96.79) | (90.06) |
| Contig | 8.33 | **7.08** | 12.00 | 11.75 | 12.45 | 10.32 |
| count | **(96.30)** | (95.52) | (91.43) | (91.78) | (94.44) | (86.02) |

**Table S5.** Wilcoxon signed-rank test results (p-values) for Kraken vs other methods.

|  | Centrifuge | RFPlasmid | mlplasmids | PlaScope | Platon |
| --- | --- | --- | --- | --- | --- |
| **Sensitivity** |  |  |  |  |  |
| Contig length | 1.000 | **<0.001** | 1.000 | **<0.001** | **<0.001** |
| Contig count | 1.000 | **0.00350** | **<0.001** | **<0.001** | **<0.001** |
| **Specificity** |  |  |  |  |  |
| Contig length | 0.137 | **<0.001** | **<0.001** | 1.000 | 1.000 |
| Contig count | 0.488 | **0.00327** | **<0.001** | 1.000 | 1.000 |
| **Precision** |  |  |  |  |  |
| Contig length | 0.122 | **<0.001** | **<0.001** | 0.527 | 1.000 |
| Contig count | 0.488 | **0.00430** | **<0.001** | 1.000 | 1.000 |
| **Negative predictive value** |  |  |  |  |  |
| Contig length | 1.000 | **<0.001** | 1.000 | **<0.001** | **<0.001** |
| Contig count | 1.000 | **0.0262** | **0.00468** | **<0.001** | **<0.001** |
| **Accuracy** |  |  |  |  |  |
| Contig length | 0.458 | **<0.001** | **<0.001** | **<0.001** | **<0.001** |
| Contig count | 1.000 | **0.00191** | **<0.001** | **<0.001** | **<0.001** |
| **F1-score** |  |  |  |  |  |
| Contig length | 0.495 | **<0.001** | **<0.001** | **<0.001** | **<0.001** |
| Contig count | 1.000 | **<0.001** | **<0.001** | **<0.001** | **<0.001** |

Wilcoxon signed-rank test was performed for each metric using values shown in **Figure 1** and **Figure S3**.
Bonferroni correction was applied across each row. p-values < 0.05 are shown in bold.

**Table S6.**
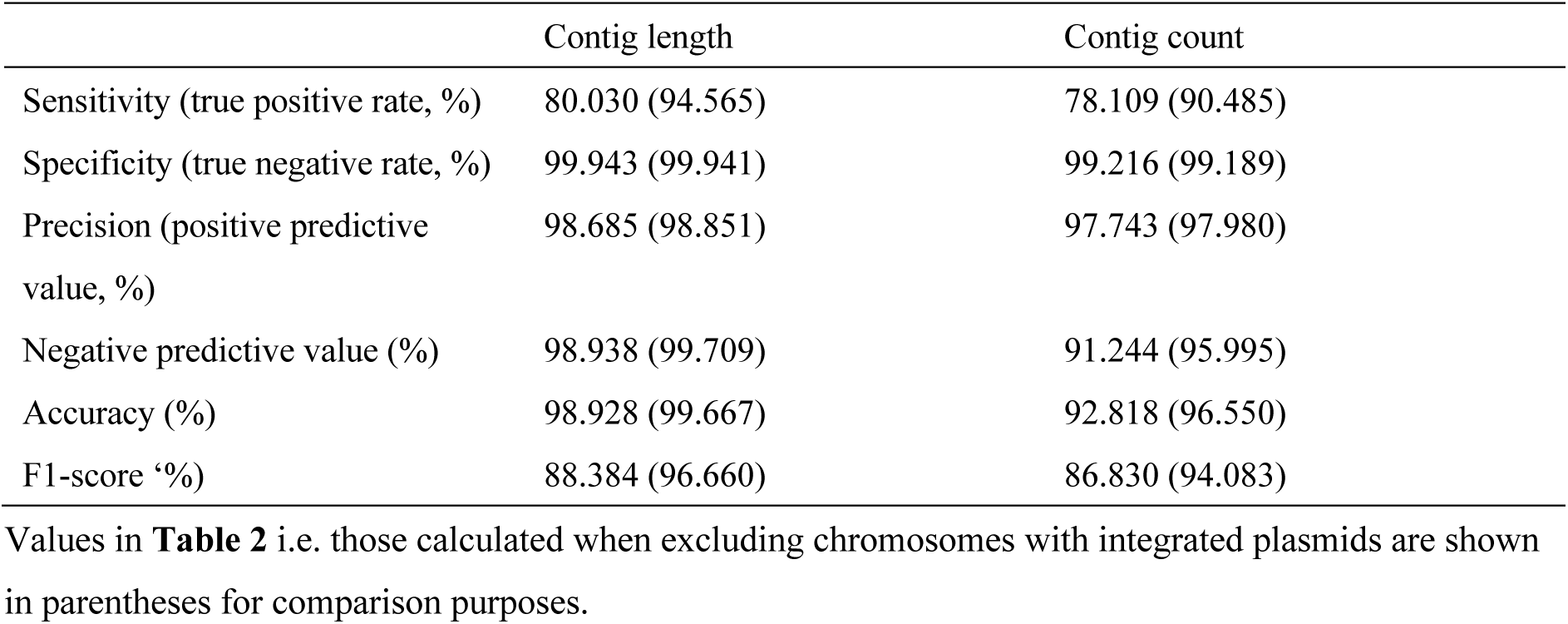
Performance of Centrifuge using the database created without excluding chromosomal sequences containing integrated plasmid sequences.

|  | Contig length | Contig count |
| --- | --- | --- |
| Sensitivity (true positive rate, %) | 80.030 (94.565) | 78.109 (90.485) |
| Specificity (true negative rate, %) | 99.943 (99.941) | 99.216 (99.189) |
| Precision (positive predictive value, %) | 98.685 (98.851) | 97.743 (97.980) |
| Negative predictive value (%) | 98.938 (99.709) | 91.244 (95.995) |
| Accuracy (%) | 98.928 (99.667) | 92.818 (96.550) |
| F1-score (%) | 88.384 (96.660) | 86.830 (94.083) |

Values in **Table 2** i.e. those calculated when excluding chromosomes with integrated plasmids are shown in parentheses for comparison purposes.

**Table S7.** Performance of mlplasmids for *K. pnuemoniae* genomes (n=69).

|  | Contig length | Contig count |
| --- | --- | --- |
| Sensitivity (true positive rate, %) | 96.158 | 95.761 |
| Specificity (true negative rate, %) | 99.771 | 93.032 |
| Precision (positive predictive value, %) | 95.770 | 86.054 |
| Negative predictive value (%) | 99.792 | 97.995 |
| Accuracy (%) | 99.585 | 93.878 |
| F1-score (%) | 95.963 | 90.649 |

**Table S8.** mlplasmids performance with a requirement of a minimum posterior probability of 0.7 for classification.

|  | Contig length | Contig count |
| --- | --- | --- |
| Sensitivity (true positive rate, %) | 90.241 | 78.794 |
| Specificity (true negative rate, %) | 99.882 | 98.323 |
| Precision (positive predictive value, %) | 97.618 | 95.335 |
| Negative predictive value (%) | 99.478 | 91.424 |
| Accuracy (%) | 99.390 | 92.403 |
| F1-score (%) | 93.785 | 86.279 |

**Table S9.** Performance metrics of Kraken in the cross validation analysis.

|  | Group A | Group B | Group C | Overall | Base database* |
| --- | --- | --- | --- | --- | --- |
| **Sensitivity (true positive rate, %)** |  |  |  |  |  |
| Contig length | 96.699 | 97.107 | 98.861 | 97.547 | 96.507 |
| Contig count | 89.984 | 89.883 | 92.784 | 90.796 | 90.796 |
| **Specificity (true negative rate, %)** |  |  |  |  |  |
| Contig length | 99.952 | 99.996 | 99.975 | 99.975 | 99.973 |
| Contig count | 99.584 | 99.718 | 99.497 | 99.594 | 99.378 |
| **Precision (positive predictive value, %)** |  |  |  |  |  |
| Contig length | 99.103 | 99.934 | 99.505 | 99.517 | 99.472 |
| Contig count | 98.917 | 99.355 | 98.684 | 98.983 | 98.449 |
| **Negative predictive value (%)** |  |  |  |  |  |
| Contig length | 99.820 | 99.839 | 99.942 | 99.868 | 99.813 |
| Contig count | 95.922 | 95.320 | 97.138 | 96.136 | 96.128 |
| **Accuracy (%)** |  |  |  |  |  |
| Contig length | 99.784 | 99.844 | 99.921 | 99.851 | 99.796 |
| Contig count | 96.732 | 96.510 | 97.558 | 96.927 | 96.777 |
| **F1-score (%)** |  |  |  |  |  |
| Contig length | 97.886 | 98.500 | 99.182 | 98.522 | 97.967 |
| Contig count | 94.239 | 94.382 | 95.643 | 94.713 | 94.468 |

\*Values in this column are from **Table 2** and shown for comparison purposes.

**Table S10.** See the excel file.

## Supplementary Methods

### Commands used for running Kraken

#### 1. Download the NCBI taxonomy

kraken-build ╌download-taxonomy ╌db database

#### 2. Add genomes to the library

for file in reference/*.fasta; do

kraken-build ╌add-to-library ${file} ╌db database

done

#Note that taxonomy information was added to the headers of fasta files. For example, the fasta file of *K. pneumoniae* chromosome NZ_CP026130.1 begins with “>NZ_CP026130.1|kraken:taxid|573”, the fasta file of *K. variicola* subsp. *variicola* chromosome NZ_CP020847.1 begins with “>NZ_CP020847.1|kraken:taxid|2590157”, and the fasta file of plasmid AP014611 begins with “>AP014611.1|kraken:taxid|36549”.

#### 3. Build the database

kraken-build ╌build ╌db database

#### 4. Classify sequences

for file in assemblies/*.fasta; do

kraken ╌preload ╌db database ${file} > $(basename ${file/.fasta})_output.txt done

### Commands used for running Centrifuge

#### 1. Build the database

centrifuge-build −p 10 ╌conversion-table seqid_to_taxid.map ╌taxonomy-tree nodes.dmp ╌name-table names.dmp database.fna chromosome_plasmid_db

#Note that the corresponding line in the nodes.dmp file was modified as follows: from

36549 | 28384 | no rank | | 0 | 0 | 11 | 1 | 0 | 1 | 0 | 0 | |

to

36549 | 28384 | species | | 0 | 0 | 11 | 1 | 0 | 1 | 0 | 0 | |

#### 2. Classify sequences

for file in assemblies/*.fasta; do

centrifuge −f −p 10 ╌reorder −x chromosome_plasmid_db −U ${file} −k 1 ╌report-file $(basename ${file/.fasta})_summary.txt −S $(basename ${file/.fasta})_output.txt done

### Commands used for running RFPlasmid

python3 rfplasmid.py ╌species Enterobacteriaceae ╌input assemblies/ ╌jelly ╌threads 8 ╌out output_directory

### R commands used for running mlplasmids

plasmid_classification(path_input_file = fasta_file, full_output = TRUE, species = “Klebsiella pneumoniae”)

### Commands used for running PlaScope

plaScope.sh ╌fasta my_fastafile.fasta −o output_directory ╌db_dir path/to/DB ╌db_name Klebsiella_PlaScope ╌sample name_of_my_sample

### Commands used for running Platon

platon ╌db path/to/DB ╌threads 10 ╌output results/ my_fastafile.fasta

